# Optimal Prescriptive Treatments for Ovarian Cancer with Genetic Data

**DOI:** 10.1101/2024.10.08.617334

**Authors:** Alkiviadis Mertzios, Matea Gjika, Xidan Xu, Samayita Guha, Neelkanth M. Bardhan, Subodha Kumar, Angela Belcher, Georgia Perakis

## Abstract

Among the cancers affecting the female population, Ovarian Cancer (OC), while being relatively rare, is the leading cause of gynecological cancer-related deaths, with overall 5-year survival rates of approximately 50% for all stages combined. This is because of the challenges associated with the diagnosis, resulting in detection at advanced stages of OC, coupled with the slow progress in effective treatment options since the approval of platinum-based chemotherapy in the late 1970s. There has been a relative lack of sophisticated methods based on Machine Learning (ML) models that use genetic data for better prediction of Ovarian Cancer outcomes and result in more effective treatment recommendations. Therefore, there is an unmet clinical need to create models that allow physicians to make informed decisions based on all available data, including patient demographic, social, health, and genomic data. Hence, we develop new techniques for leveraging genetic information in prescribing optimal treatments for patients with OC, using a publicly available dataset from the Prostate, Lung, Colorectal and Ovarian Cancer (PLCO) trial. Our approach is able to transform genotype sequencing information into a simple tabular form that can then be used as the input to any ML model. Coupled with the recorded treatment regimen and clinical parameters of matched patients from the genetic dataset, we estimate the treatment effect in terms of mortality prediction and use it to prescribe the optimal treatment for any given patient. By including the genetic features engineered through our proposed method, our models have a higher accuracy than the models without genetic information embedded. The increase in predictive accuracy demonstrates the improved efficacy of our method in the predictive setting. Furthermore, in the prescriptive setting, the models including genetic features output different treatment choices for patients, showing the impact of their inclusion. This is further highlighted by the feature importance of the genetic features such as mutations in the FAT3, BRCA1, BRCA2, and NF1 genes, where they rank highly with a tighter aggregation of the top features, relative to the sharp drop-off in feature importance after the top feature in the models without genetic data. Taken together, in summary, our models will allow oncologists to make more informed and accurate decisions, incorporating a patient’s genetic data with all other available clinical information, which has the potential for improved prognosis and better long-term survival outcomes for Ovarian Cancer patients.

## 1. Introduction

Ovarian Cancer (OC) is a prominent health concern for women around the world, being the third most common gynecologic malignancy. According to the GLOBOCAN 2020 estimates of the global cancer burden (Sung et al., 2021), globally, there were approximately 314, 000 cases of ovarian cancer, resulting in over 200,000 deaths related in 2020 alone (Cabasag et al., 2022). The same study found an incidence rate of 4.2% for OC around the globe. In the United States, the incidence rate for OC was 1.3% for the years between 2012-2014 for those with no other forms of cancer, while the mortality rate was 6.7 in 100,000 (total population) in 2015 (Torre et al., 2018). Furthermore, OC has a relatively poor long-term prognosis, with the 5-year survival of advanced-stage OC patients is approximately 20% (Huang et al., 2022). This makes OC a major global health problem, for which new methods have to be developed in order to deal with it in an effective manner.

Till date, some Machine Learning (ML) approaches have been employed in the OC problem. The existing literature focuses on the problem of prediction of OC, meaning given a person’s certain attributes the ML model outputs the probability that OC will develop in the future. For example, Lu et al. (2020) use a decision tree to predict OC incidence, while also identifying features of importance. In this work, it was reported that human epididymis protein 4 (HE4) and carcinoembryonic antigen (CEA) had the most predictive power. On the other hand, Paik et al. (2019) use Gradient Boosting to predict OC mortality in patients diagnosed with Epithelial Ovarian Cancer.

While these approaches are successful in predicting OC, they are only informative in the sense that the patient knows if she will survive or not, with no impact on the decision-making process for the patient, or the generation of actionable insights for the doctors. To have a model that helps in the decision-making, it is desirable to develop a model that recommends a treatment plan that maximizes the probability of a patient’s survival, given the patient’s characteristics.

Another aspect of OC that seems to elude the literature, is the use of genetic data. Prior family history of breast, uterine, or ovarian cancer can increase risk of future OC (Mori et al., 1988). Other genetic factors such as the presence of germline mutations can also contribute to increased risk of OC. For example, mutations in the genes BRCA1, BRCA2 and TP53 are linked to an increased probability of developing the disease (Ford and Easton, 1995; Boyd and Rubin, 1997).

In this work, we reconcile these two bodies of work and develop a method for evaluating the extent of a gene’s mutation. We use these new genetic features to train a causal model that allows us to assess the effect of each treatment in the population, allowing us to make accurate and effective prescriptions of individualized treatments. To the best of our knowledge, this is the first work that uses genetic information for prescription of the optimal treatment for a given patient diagnosed with OC.

We note that our method for engineering genetic features leads to improved predictions both in mortality as well as decision of treatment. Inclusion of genetic data not only improves the accuracy of predictions, but also genetic features rank high in feature importance within our proposed models. This showcases the efficacy of our approach, which we contrast with models that do not include genomic data. In the predictive setting, incorporating genetic data improves accuracy of prediction, while feature importance of genes such as “FAT3” ranks high among the features. In the prescriptive setting, we enjoy similar improvement, with genetic data offering different treatments to patients, thus showing the impact of including genetic data with our proposed method. Again, genetic features rank high in importance out of all features that were included in the model.

Our ML approach allows us to succinctly represent genetic data of ovarian cancer patients, and use them to prescribe treatments and make predictions about a patient’s mortality. Also, we provide a ranking method for choosing the best treatment for a given patient using the treatment effect that we learn for every treatment. To the best of our knowledge, this is the first approach that uses ML to extract information from the patients genome, and outputs an aggregate for each gene. We are able to do this by taking the CADD score (Rentzsch et al., 2018; Schubach et al., 2024) of each Single Nucleotid Polymorphism (SNP) (Wright, 2005) and aggregating. The novelty of our approach is combining this data with causal models (Wager and Athey, 2018; Li et al., 2017) to prescribe individual treatments using the patients’ genetic information.

## 2. Related Work

Causal inference has seen significant growth in recent years, driven in part by advancements in machine learning techniques. A wide range of methods exists for estimating causal effects, which refer to the difference an intervention or treatment makes in outcomes. This process is often referred to as counterfactual estimation (Morgan and Winship, 2014), as we cannot observe both the treated and untreated outcomes for the same individual. To address this, researchers often rely on re-weighting methods, which assign weights to treatment effects based on their prevalence, or on stratification and matching techniques, which group similar individuals to more accurately estimate the treatment’s impact on outcomes (Yao et al., 2021).

There is also a large body of work dedicated to extending classical ML models to the causal inference setting (Wager and Athey, 2018; Li et al., 2017; Bakhitov and Singh, 2022). An example of this is causal decision trees (Athey and Imbens, 2016), which adapt the well known decision tree classifier to the causal inference paradigm. Therein, the authors use an honest approach to training; the samples that are used for creating the tree are not reused for estimation of treatment effects. This allows for a predictor with unbiased estimates of the treatment effect. Furthermore, in Athey et al. (2019), the authors further generalize the random forest framework, by offering a method for non-parametric statistical estimation based on random forests.

On the healthcare side of things, there has been a lot of work in personalized treatments for patients. In Chen et al. (2021), the authors provide treatment prescriptions to patients with early stage breast cancer patients undergoing radiotherapy. The authors are able to achieve a reduced risk of tumor recurrence, while minimizing the side effects caused by radiation treatment.

In OC, machine learning methods are applied to a variety of tasks, from OC diagnosis to estimation of drug responses in patients. In Yue et al. (2021), the authors leverage the power of neural networks to diagnose OC, using plasma related data. Another task that is very relevant to the OC is to decide which drug would help a patient the most. In this context, machine learning can be used to accurately and efficiently predict the effect of certain drugs and ultimately provide doctors with recommendations for which drug prescription to follow (Huang et al., 2018).

Finally, genetic mutations have long been associated with OC (Ford and Easton, 1995; Boyd and Rubin, 1997). One such case is mutations that affect genes regulating the homologous recombination mechanism, leading to a state called Homologous Recombination Deficiency (HRD) (da Cunha Colombo Bonadio et al., 2018). Patients with HRD are unable to perform double strand repair of DNA. However, this allows for the use of a class of drugs called poly ADP ribose polymerase (PARP) inhibitors, which cause synthetic lethality to cancer cells (Konstantinopoulos et al., 2015).

We extend this body of work by introducing novel methods for including genetic data in ML models. Specifically, we develop a method for succinct representation of genetic data that we combine with demographic and cancer data of each patient. This allows us to get more accurate models that use genetic data efficiently. We develop a predictive model that can help physicians and patients make more accurate assessments of their health situation, and also a prescriptive model that can help oncologists make genetic informed decisions for the treatment of OC patients, thus increasing survivability.

## 3. Method

A schematic showing the workflow used in our methods is depicted in Figure 1. We describe the methods used to succinctly extract genetic information from the patients. For the present work, we use data from the Ovarian Cancer cohort, collected in the Prostate, Lung, Colorectal, and Ovarian Cancer Screening Trial (PLCO) (Prorok et al., 2000). In this clinical trial, subjects where randomly assigned into one of two groups, where one group enjoyed yearly diagnostic tests related to one of the aforementioned four cancer types. The goal of the trial was to investigate if such regular screening is able to reduce mortality, if cancer was to develop in the screened population. Along with the diagnostic tests, demographic and genetic data were collected, as well as staging, histological, grading and treatment data for those individuals diagnosed with cancer. This makes the PLCO dataset ideal for investigating relations between genetic data and treatment, with the ultimate goal being prescribing the optimal treatment.

**Figure 1.**
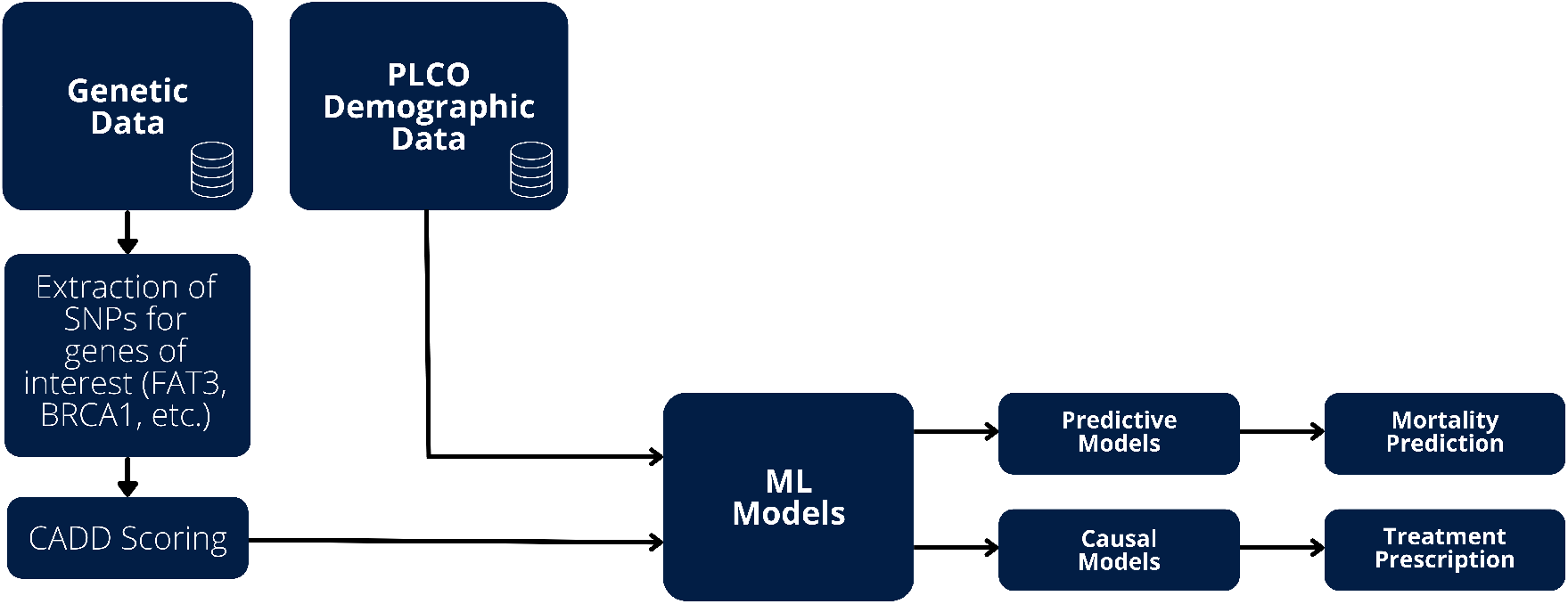
Schematic of the workflow of our method, showing the different pipelines for handling genetic *v*. demographic data, which are then fed into our ML models. Combining the genetic and demographic data sources leads to higher accuracy and better results for both mortality prediction and treatment prescription.

The genetic data in the PLCO dataset are in the form of Single Nucleotide Polymorphisms (SNPs). The patients were genotyped using whole genome high-density genotyping arrays, specifically Illumina (Zhao et al., 2017). Genotypes of subjects are collected for a set of predetermined genome positions. Quality control methods are also applied to ensure the reliability of the data.

The genetic information we described are in the form of large files, and cannot be used directly in ML models due to the sheer size and format of the data. Each gene may contain hundreds of SNPs, making it infeasible to define a feature for each one. In addition, different subjects are genotyped in different arrays and as such might not contain the same set of SNPs as each other. Consequently, in order to efficiently use the genetic data, we need to leverage a method that succinctly represents the genetic state of each subject with few features per gene.

In order to achieve this, we use the Combined Annotation-Dependent Depletion (CADD) score (Rentzsch et al., 2018). CADD is a dataset that calculates a probability of deleteriousness for each mutation in the human genome. The authors achieve this by training a logistic regression model that takes as inputs different mutations, ones that have survived through the evolution process and are likely to be innocuous, and simulated ones that are likely to be deleterious. Then these are annotated using their genetic context and properties and the model is trained. The result is a probability of deleteriousness for each given mutation. Our method takes the values produced by CADD and computes the PHRED scores (Ewing et al., 1998; Ewing and Green, 1998). The PHRED scores are computed by taking the logarithm of the percentiles of each CADD score extracted. As such, they give a regularized quantity for each mutation, with higher scores relating to a greater chance of deleteriousness. We aggregate the PHRED scores by summing all of the scores within each gene, and also noting the number of heterozygous and homozygous mutations. Then, the number of mutations along with the summed and mean PHRED scores are gathered for each gene of interest and used as features in our ML models.

This study applies Causal Forest methodologies to elucidate the diverse effects of treatments effects. Concurrently, predictive models are deployed to forecast mortality outcomes for patients with Ovarian cancer, empowering healthcare professionals with foresight into potential outcomes.

Within the predictive models employed, the key outcome variable f dthovar is a dichotomous indicator of mortality, with a value of 1 signifying death due to ovarian cancer and 0 representing all other causes (Prorok et al., 2000). We utilize a comprehensive array of patient characteristics alongside genomic data to analyze mortality risk. Specifically, we want to investigate the contributory role of genomic information in enhancing the models’ performance. In addition to genomic insights, we examine alternative methodologies, such as various encoding techniques, to improve the interpretability of our models. Further details on the predictive modeling approach and variable specifications are elaborated in the subsequent section.

## 4. Dataset

The PLCO cancer screening trial (Prorok et al., 2000) collected data for 155,000 participants. Participants were enrolled between November 1993 and July 2001. Data were collected on cancer diagnoses through 2009 and mortality through 2018. For Ovarian Cancer, the number of participants was 78, 209. Among factors excluding participants from the trial were age (subjects younger than 60 or older than 74), history of cancer for the cancer of interest, removal of both ovaries (for participants before October 1996), participants of other cancer trials and patients previously treated with Tamoxifen or Evista/Raloxifene in the 6 months prior to randomization.

The predictive models examine a carefully defined subset of the participants described above. Selection criteria are established to identify individuals with baseline ovarian presence. Furthermore, participants are required to be part of the overall Genome-Wide Association Study (GWAS) cohort, which denotes individuals with genomic data recorded. Additionally, only those with a confirmed cancer diagnosis are selected for analysis. These stringent selection parameters culminated in the formation of a targeted cohort of 302 patients.

### 4.1. Feature Choices

In our predictive modeling framework, we capture a spectrum of 65 distinct features per patient. These features include a set of 35 non-genomic variables (Table 2), and an additional 30 genomic features derived from key genes implicated in cancer pathology: NF1, TP53, FAT3, BRCA1, and BRCA2 (Korf, 2000; Olivier et al., 2010; Guo et al., 2021; Singh et al., 2022; Petrucelli et al., 2023) respectively. For these genomic markers, we calculate the mutation count, the mean, and aggregated PHRED score for each individual.

**Table 1:** Accuracy and AUC values for Logistic Regression.

|  | Accuracy | AUC |
| --- | --- | --- |
| Model trained w/o genomic data | 0.7541 | 0.7168 |
| Model trained w/ genomic data | 0.7705 | 0.7230 |

**Table 2:**
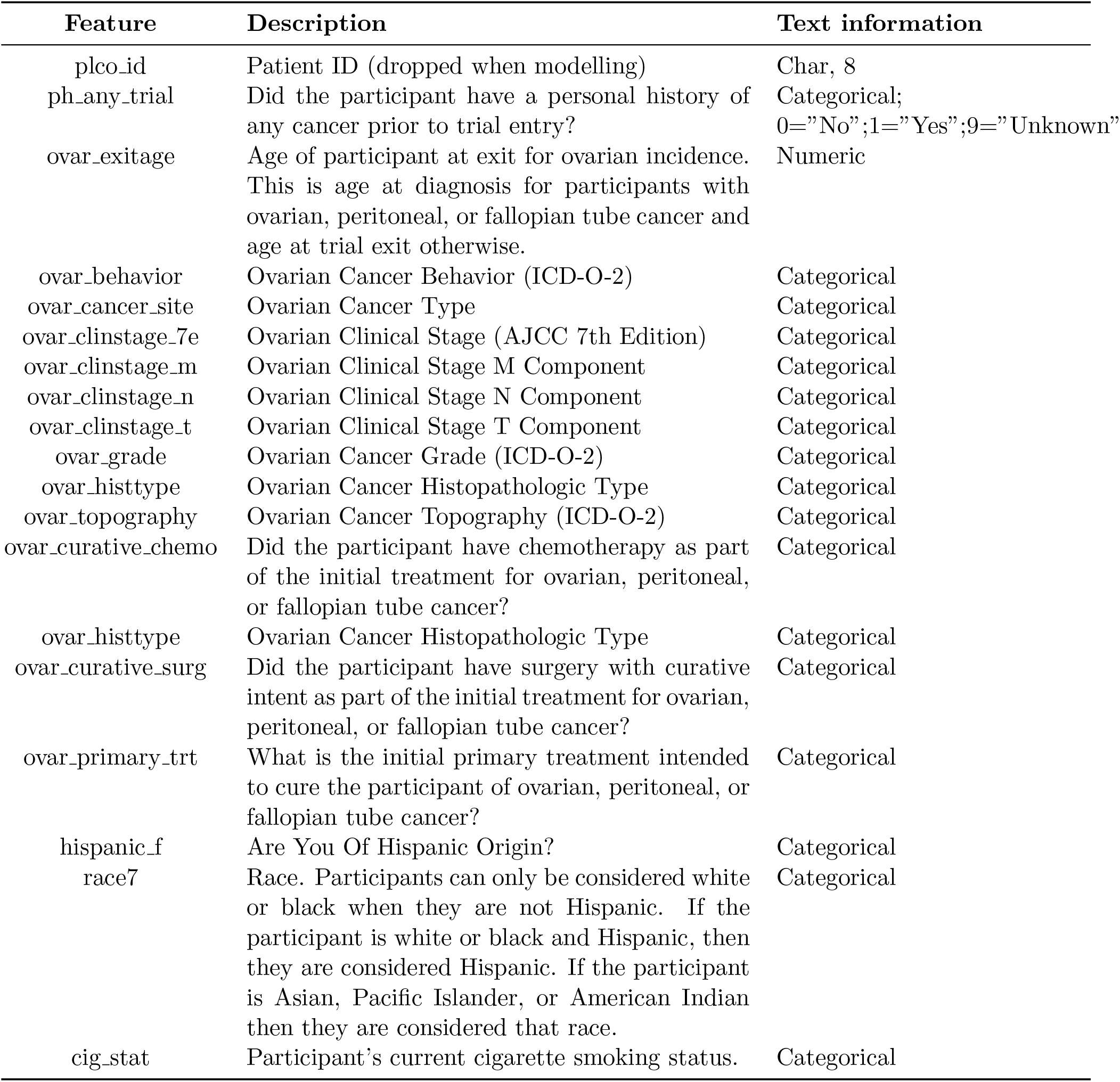
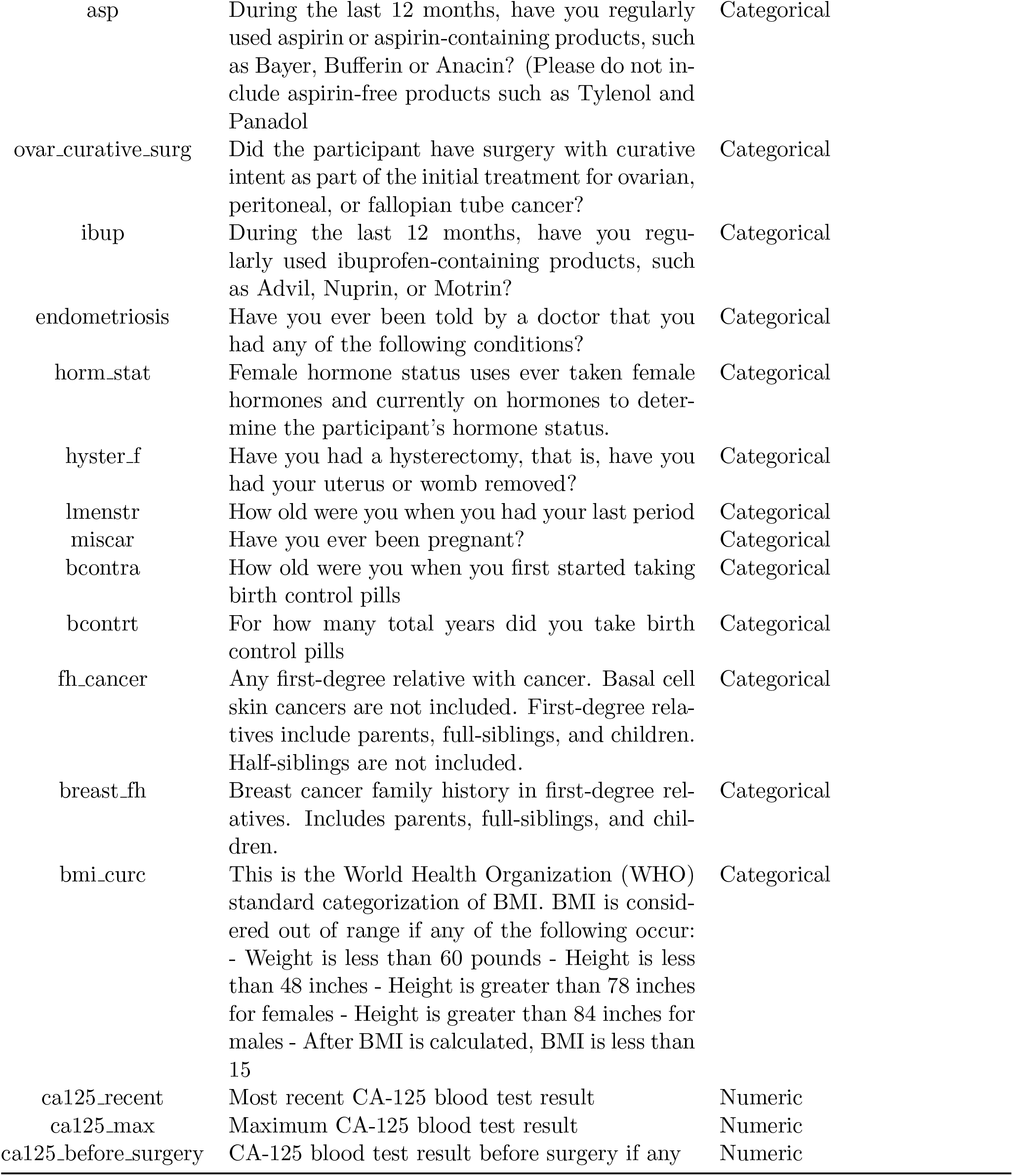
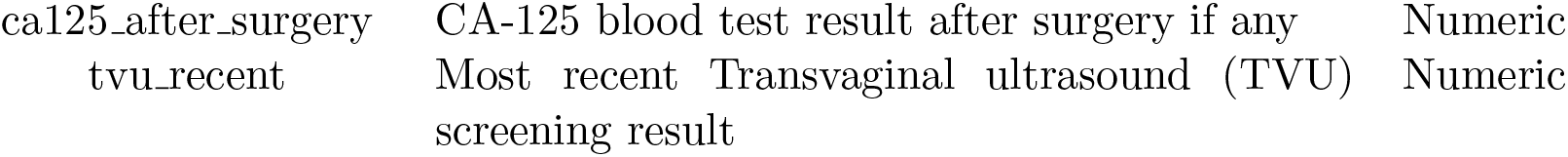
Non-genomic feature List for predictive model.

Notably, special emphasis is given to the evaluation of CA-125 levels, a key non-genomic datum, due to its established correlation with ovarian cancer (Charkhchi et al., 2020). CA-125 is monitored longitudinally, allowing for the extraction of four pivotal features: the most recent measurement, the peak value, and the comparative levels pre- and post-surgery, thereby providing a temporal profile of this critical biomarker across different stages of patient care.

### 4.2. Data Preprocessing & Missing Value Imputation

To facilitate rigorous analysis, the dataset is subjected to a series of preprocessing procedures. In the case of categorical variables, the mode imputation technique is employed for missing data, assigning the most frequent category to fill gaps.

Features exhibiting more than 50% missing data are deemed unreliable and subsequently removed, following the guidelines discussed in the literature (Acuña and Rodriguez, 2004). This 50% threshold is carefully considered since, as mentioned by Lin and Tsai (Lin and Tsai, 2020), the majority of data imputation research does not discuss data imputation for more than 50% missing data, and among the few studies that do, some conclude that data with over 50% missing entries should be excluded (Acuña and Rodriguez, 2004). Features being dropped include CA-125 level after surgery with only 10 values.

Additionally, We address the pivotal CA-125 biomarker in our study by employing imputation techniques. Specifically, we apply the mean imputation methods to preserve the integrity of this crucial variable.

To prepare categorical data for machine learning algorithms, frequency encoding is applied, which assigns to each categorical value the number of its occurrences. We consider frequency encoding for its ability to maintain interpretability, especially when assessing feature importance.

## 5. Results on Real Data

In this section, we describe our experiments on the PLCO dataset. We train models for two paradigms, the predictive and prescriptive. The predictive model allows for accurate predictions of mortality, while the prescriptive model dictates which treatment is most beneficial given each patients physical background. Both models enjoy increased accuracy as a result of our method of interpreting genetic data.

### 5.1. Mortality Predictive Model

For the predictive model, we use Logistic Regression (Tibshirani, 1996) for its interpretability. To discern the contribution of genomic data to prediction accuracy, we train two versions of the model, one with and one without the genetic data. We compare between the two and show that the inclusion of genetic data improves prediction accuracy as well as area under the curve (AUC). Besides model comparison, we also consider a naive baseline model, simply predicting the majority class. The accuracy of the naive baseline model in our work is 60.6%.

We divide the dataset into an 80% training set and a 20% test set, adhering to standard practices for testing. Furthermore, cross-validation is employed to optimize over the hyperparameters of our ML models, preventing overfitting and ensuring the generalizability of our findings.

In table 1, we show the results for classifiers when trained with and without genomic data.

While both models outperform the baseline, we observe a clear improvement in prediction power when incorporating genetic features that were engineered using our method. Including genomic data, we have an accuracy improvement of 1.6%, a number that is significant when dealing with large cohorts and decisions that affect human life.

In figures 2 and 3 we outline the coefficients of the top 12 features in our models. Genetic features are represented using numerical allele labels. A notation of “1 | 1” denotes that the individual possesses two copies of the allele labeled as ‘1’, indicating a homozygous state. In contrast, the notation “0 | 1” signifies the presence of two different alleles, ‘0’ and ‘1’, characterizing this as a heterozygous state. Here, label ‘1’ refers to the alternate, mutated allele.

**Figure 2.**
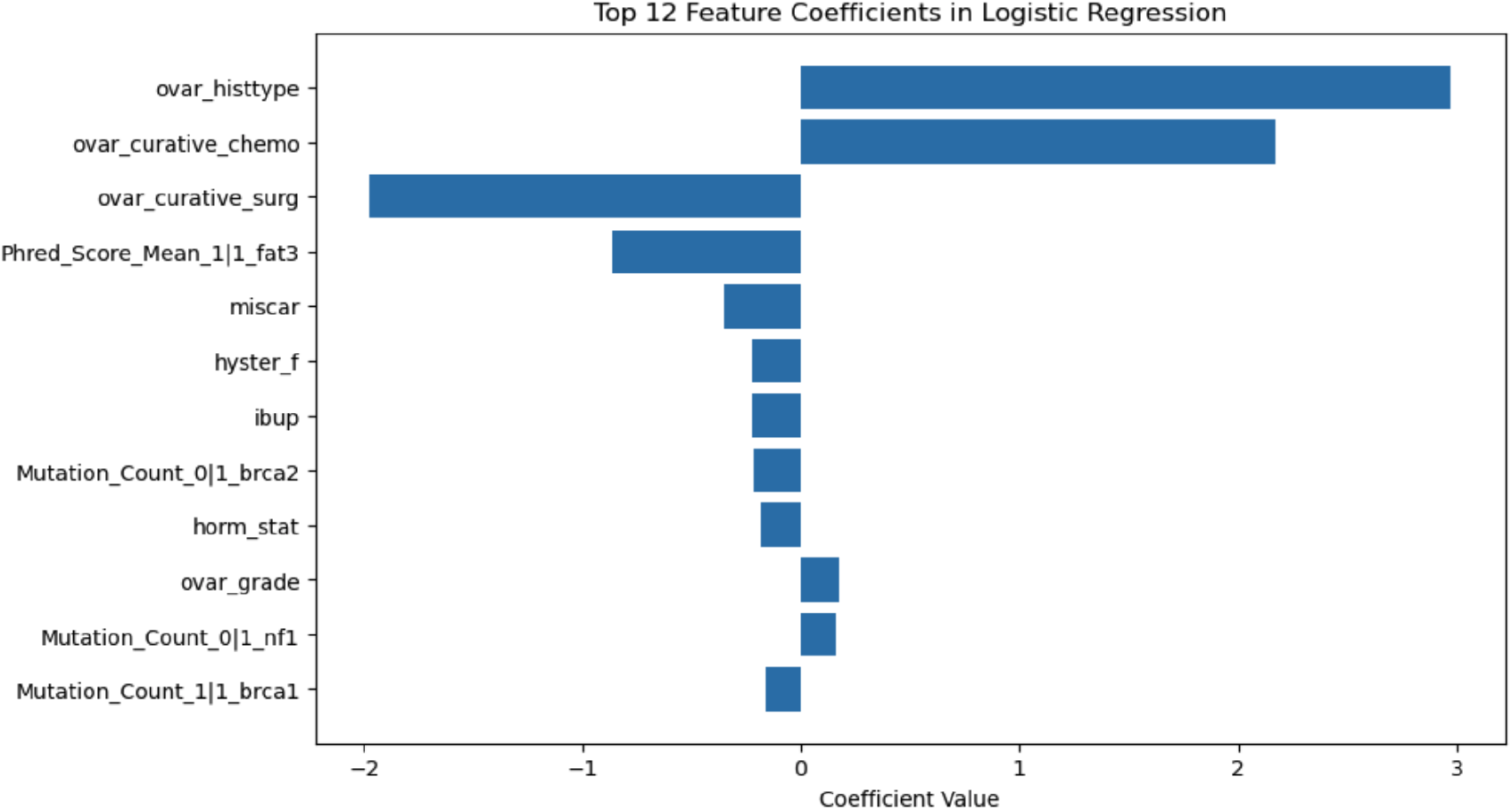
Feature Importance for Model with Genomic Data

**Figure 3.**
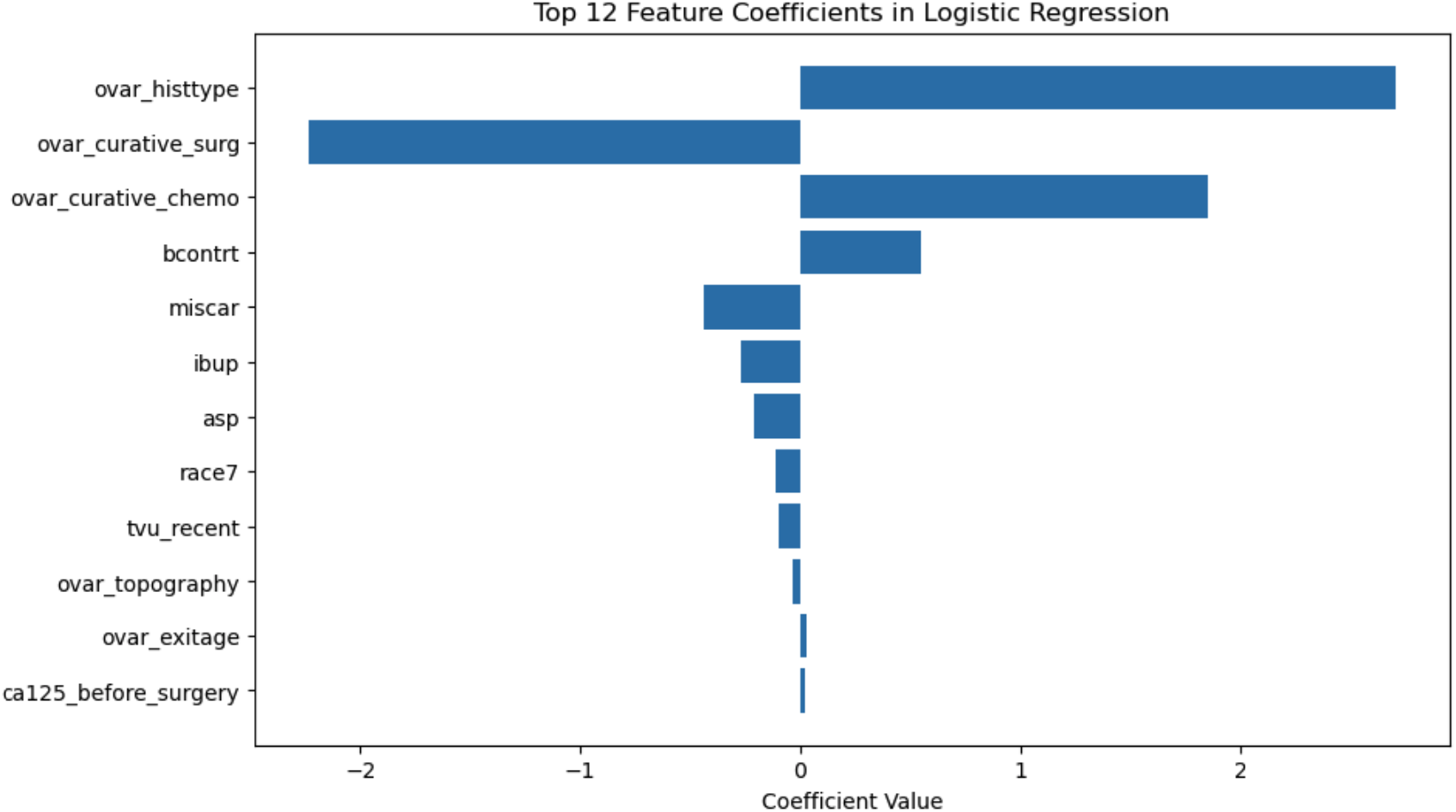
Feature Importance for Model without Genomic Data

The presence of genomic features, especially “FAT3”, within the most important features of the model including genomic data is indeed significant. While the present work uses the PLCO dataset, recent work on the tumor mutational burden from 432 OC patient samples derived from The Cancer Genome Atlas ovarian cancer data collection (TCGA-OV) independently reports FAT3 or FAT4 mutations as putative drivers of OC which are shared in 26% of early OC patients (Ban et al., 2023), thereby corroborating our results on feature importance of these genomic mutations. The prevalence of such features in our models is indicative of the potential value that genomic data holds in the context of ovarian cancer prognosis and treatment outcome prediction. Additionally, the improvement of models with genomic data and the high importance of genomic features in our model suggest these variables are potentially critical to distinguishing between mortality due to ovarian cancer versus other factors.

## 6. Causal Model

The primary objective of our causal modeling approach is to ascertain the effectiveness of individual treatments, paving the way for a tailored treatment regimen for each patient. To achieve this, we harness the power of causal forests to perform nuanced causal estimations. In our quest to understand the impact of genomic data on our analysis, we conducted a comparative study by applying our causal model to two distinct datasets: one inclusive of genomic scores and another devoid of them.

Causal forests Wager and Athey (2018) are designed to uncover and amplify the variations in treatment effects across diverse patient groups—distinguishing between those who have received the treatment and those who have not. This methodology draws on the principles of random forests, wherein causal trees are cultivated from randomly sampled data subsets and features, with the collective output of these trees providing a comprehensive estimation of treatment effects. Moreover, examining the feature importance within causal forests offers insightful revelations on the incremental value genomic data brings to our analysis.

Employing causal forests enables us to precisely estimate the causal impact of various treatments on patients within the test set. This foundational analysis informs the development of our prescriptive pipeline, which systematically identifies the optimal treatment based on the highest estimated causal effect, thereby customizing healthcare solutions to individual patients. Figure 4 shows the estimated treatment effects for different treatments. The corresponding treatment names for each code can be found in Table 3 in the appendix.

**Table 3:**
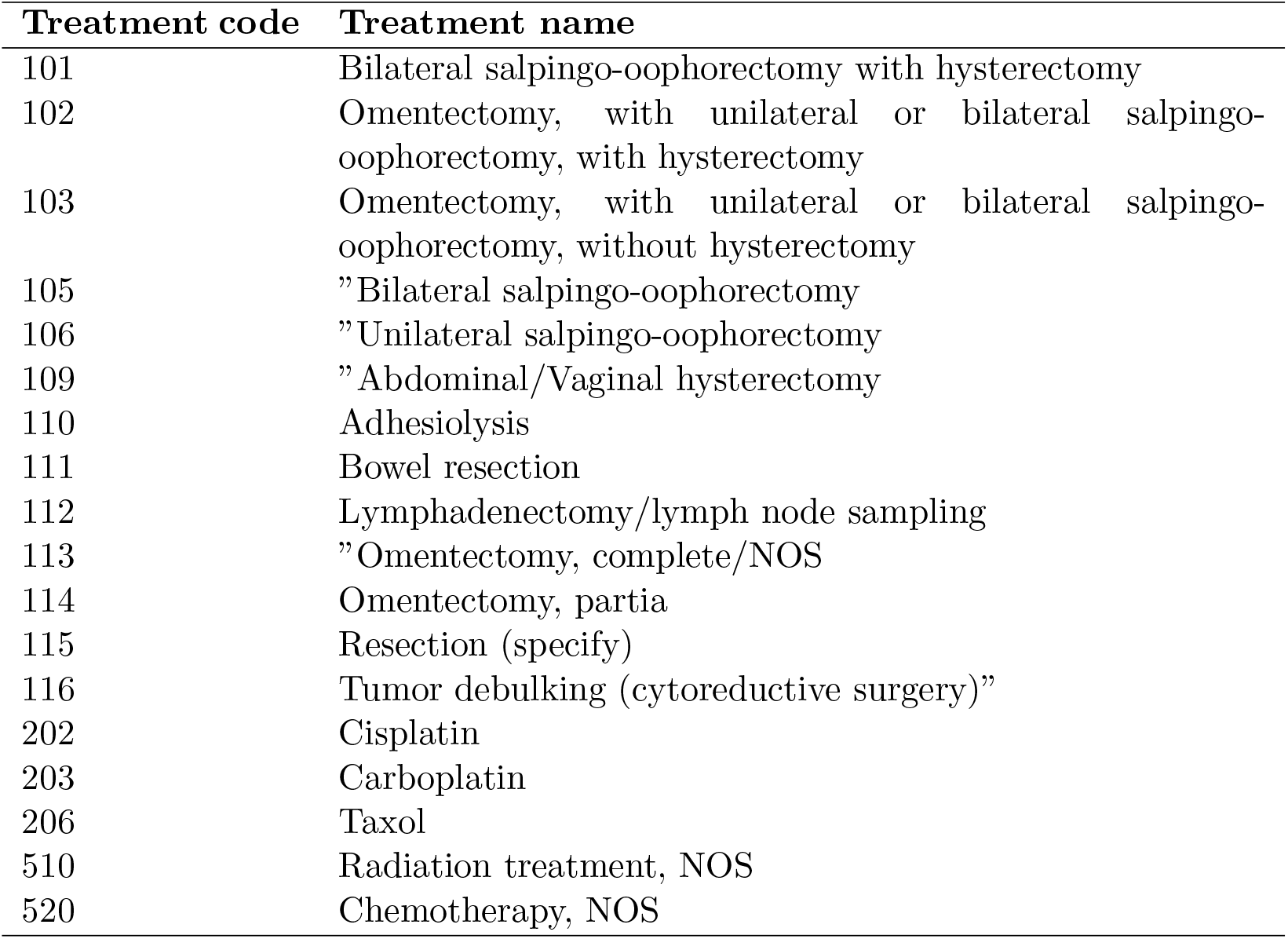
Treatment codes and names.

| **Treatment code** | **Treatment name** |
| --- | --- |
| 101 | Bilateral salpingo-oophorectomy with hysterectomy |
| 102 | Omentectomy, with unilateral or bilateral salpingo-oophorectomy, with hysterectomy |
| 103 | Omentectomy, with unilateral or bilateral salpingo-oophorectomy, without hysterectomy |
| 105 | "Bilateral salpingo-oophorectomy |
| 106 | "Unilateral salpingo-oophorectomy |
| 109 | "Abdominal/Vaginal hysterectomy |
| 110 | Adhesiolysis |
| 111 | Bowel resection |
| 112 | Lymphadenectomy/lymph node sampling |
| 113 | "Omentectomy, complete/NOS |
| 114 | Omentectomy, partial |
| 115 | Resection (specify) |
| 116 | Tumor debulking (cytoreductive surgery)" |
| 202 | Cisplatin |
| 203 | Carboplatin |
| 206 | Taxol |
| 510 | Radiation treatment, NOS |
| 520 | Chemotherapy, NOS |

**Figure 4.**
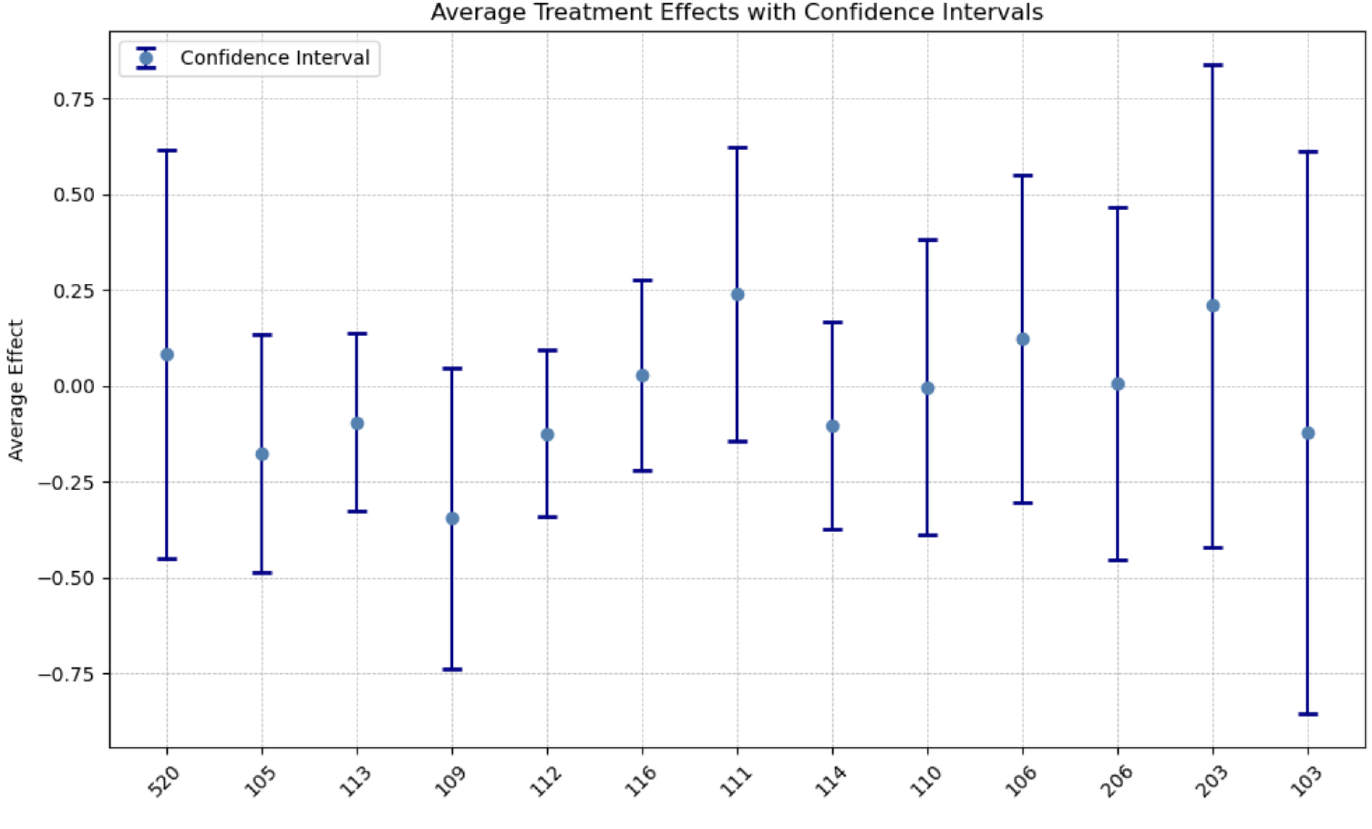
Treatment effects

Our analysis reveals a significant variation in treatment efficacy, with certain treatments showing positive outcomes and others exhibiting negative causal effects. It’s crucial to consider prescription bias in our interpretation, particularly as some treatments are more frequently administered to patients with advanced stages of cancer, potentially skewing the results.

When analysing the feature importance for both causal models, we find that for the model that includes genomic data, the most relevant variables are the age at which a patient’s ovaries were removed, followed by genomic information related to the FAT3, BRCA1, BRCA2 and NF1 genes, as seen in Figure 5. On the other hand, the causal effects in the other model are overwhelmingly dependent on the age at which ovaries were removed, with other features, such as cancer grade, smoking status, BMI and age at menopause having a significantly lower impact, found in Figure 6. This shows the genomic data provides a more granular insight into computing treatment effects for ovarian cancer.

**Figure 5.**
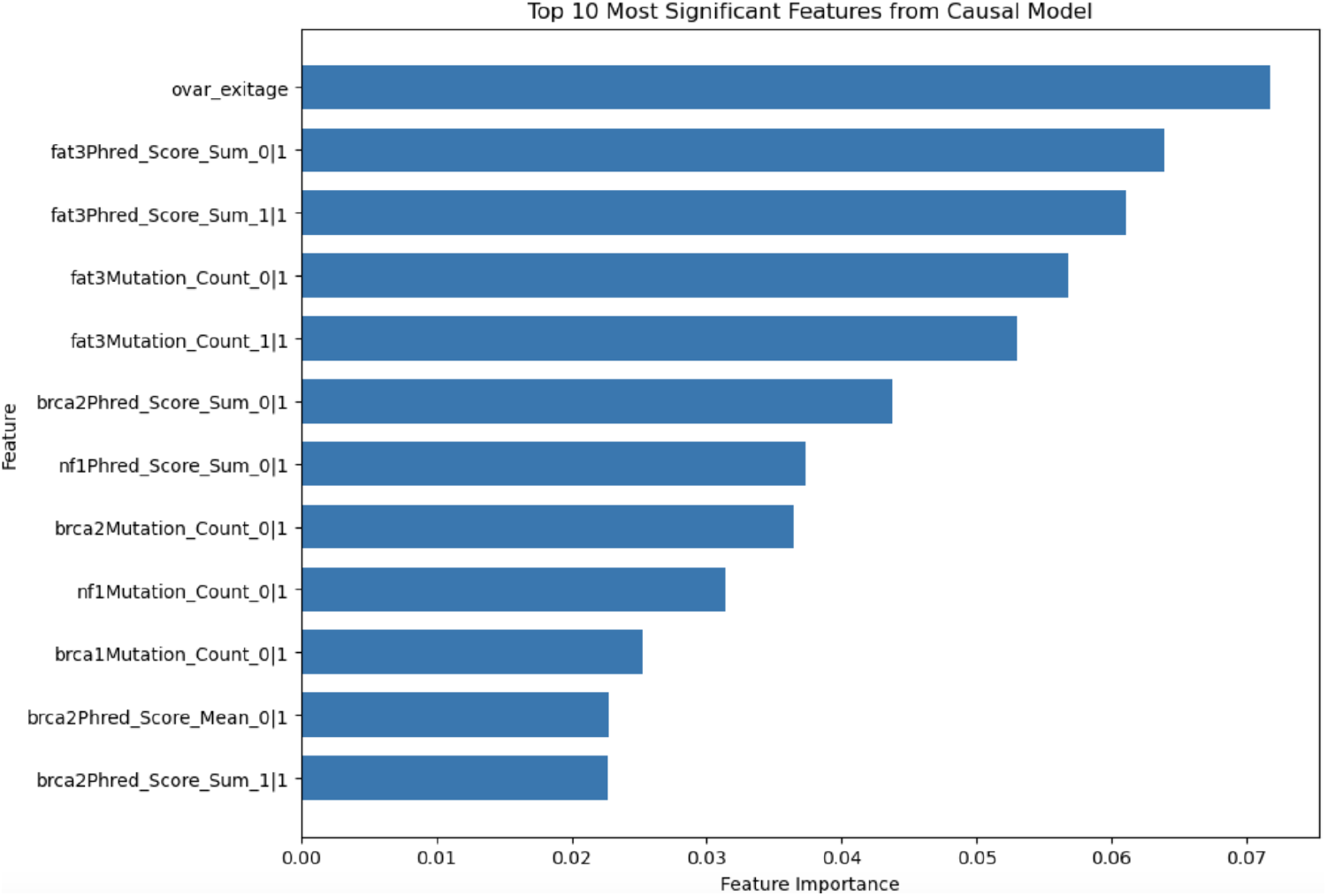
Feature importance: Causal Model with Genomic Data

**Figure 6.**
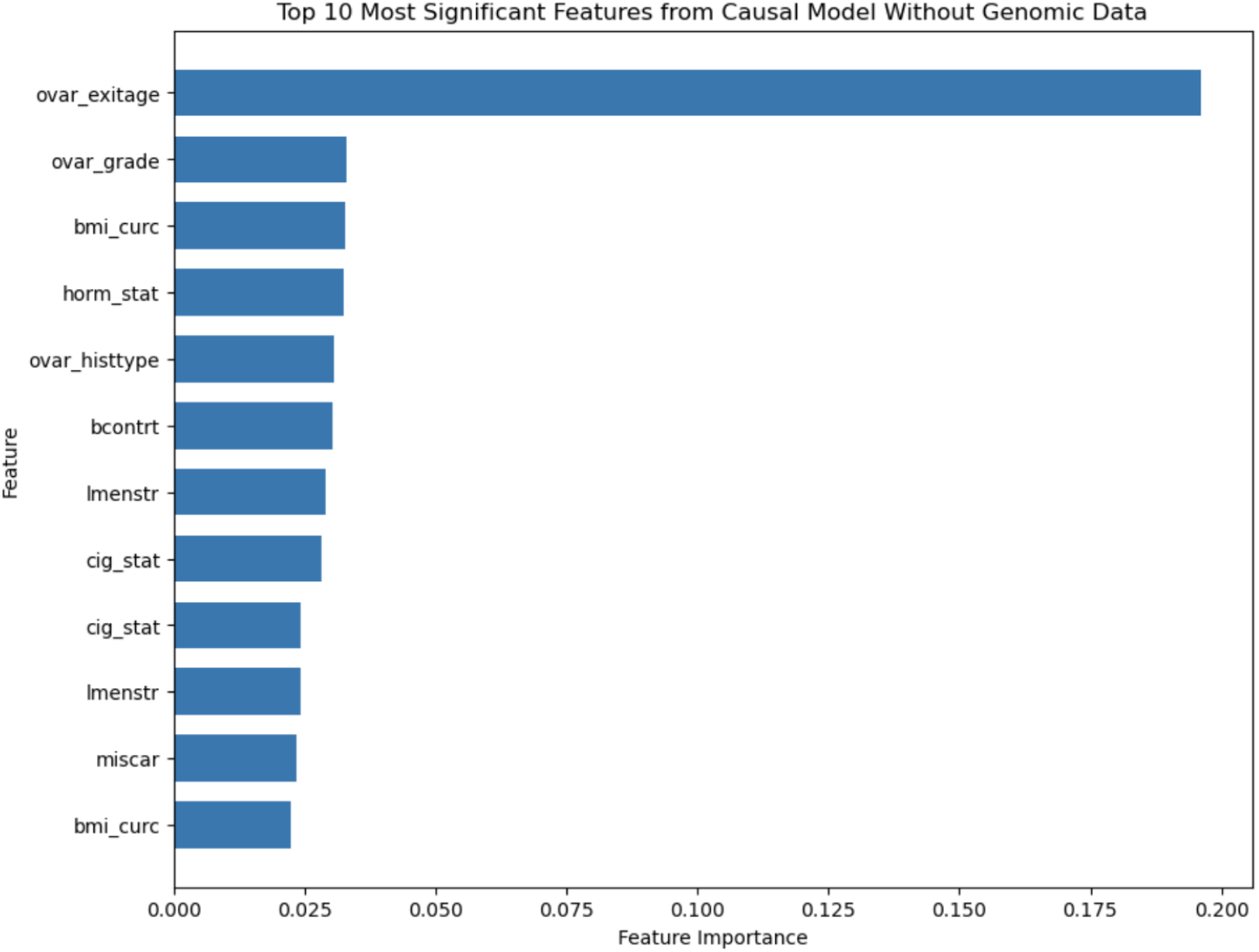
Feature importance: Causal Model without Genomic Data

Our findings are particularly striking when examining the practical implications of including genomic data in treatment selection. Among 61 patients in our test set, the use of genomic data suggests alternative treatments for 40 individuals, with 35 patients showing improved outcomes under the genomic-inclusive model. Quantitatively, the average treatment effect for the model incorporating genomic data stands at 0.33, compared to 0.32 for the model without genomic data. This marginal improvement underscores the potential of genomic data to enhance the precision of personalized treatment plans significantly.

## 7. Discussion

Given our proposed method and experiments, we see that the incorporation of genomic data in predictive as well as prescriptive models offers the model a higher capacity for accuracy. In the predictive models, the inclusion of genomic features improves accuracy as well as AUC. In addition, genomic features rank high in feature importance for the models, with the “FAT3” gene being one of the top performers.

In the causal model, inclusion of genomic features leads to different estimation of the causal/treatment effect and thus points to alternate recommendations for the treatment of patients. It is worth reiterating that the genomic features are critical in this prescription, as genomic features take the top places in feature importance among all features.

In conclusion, both the predictive and the causal models enjoy higher accuracy given the inclusion of genetic data in the training set. The predictive model can be used by patients and physicians alike to estimate the mortality risk faced by a particular patient, while the causal model can be used as a prescriptive tool to better inform treatment decisions on the part of oncologists. As such, our work offers a new tool for clinicians to incorporate the genetic data of patients in deciding optimal treatments, which could potentially lead to improved prognosis and better long-term survival of Ovarian Cancer patients.

### Limitations

While we recognize the power that our method can provide in both predictive and prescriptive models using genomic features, we note that this study is limited by the amount of available data. As mentioned previously, only 302 patients are included in the models, as those are the subjects who have been diagnosed with ovarian cancer over the duration of the PLCO trial for whom genetic data is available. As PLCO was a randomized screening trial with subjects who did not necessarily have cancer, only a very small percentage of the trial participants ended up with confirmed diagnoses of OC. This leads us to believe that a larger dataset including more cancer patients might be more ideal. However, large, comprehensive datasets containing matched treatment, genetic and demographic data for cancer patients only are very rare, especially when one focuses on a specific type of cancer such as Ovarian Cancer.

## Appendix A.

Further results in our experiments, as well as a detailed description of all features included in our models.

## Notes

### Competing Interest Statement

The authors have declared no competing interest.

